# Enhanced immune evasion of SARS-CoV-2 KP.3.1.1 and XEC through NTD glycosylation

**DOI:** 10.1101/2024.10.23.619754

**Authors:** Jingyi Liu, Yuanling Yu, Fanchong Jian, Sijie Yang, Weiliang Song, Peng Wang, Lingling Yu, Fei Shao, Yunlong Cao

## Abstract

KP.3.1.1 has surpassed KP.3 to become the new globally dominant strain, while XEC, a recombinant variant of KS.1.1/KP.3.3, is rapidly expanding across Europe and North America. Notably, both variants carry mutations, S31del of KP.3.1.1 and T22N of XEC, that could introduce new N-linked glycans on the Spike N-terminal domain (NTD), emphasizing the urgent need to assess their potential changes in viral characteristics. Here, we found that both KP.3.1.1 and XEC maintained the high ACE2-Spike binding affinity and pseudovirus infectivity of KP.3. Importantly, compared to KP.3, KP.3.1.1, and especially XEC, could further evade the neutralizing antibodies in convalescent plasma, even those elicited by KP.2-like breakthrough infections. Interestingly, both variants demonstrated increased resistance against monoclonal neutralizing antibodies targeting various epitopes on the receptor-binding domain (RBD). These suggest that the additional NTD glycosylation of KP.3.1.1 and XEC could enhance immune evasion via allosteric effects, and supports the future prevalence of XEC.

## Main

KP.3, a subvariant of JN.1, has rapidly emerged as the dominant strain in several countries and has been designated as a Variant Under Monitoring (VUM). Previous studies indicate that the unique Q493E substitution in KP.3 enhances its binding affinity for the ACE2 receptor and immuno-evasive capabilities, enabling it to outcompete KP.2 in prevalence^1-7^. Notably, KP.3.1.1, which possesses an additional S31del mutation compared to KP.3, has surpassed KP.3 to become the new globally dominant strain^8^ (Figures A and B). While KP.3.1.1 is prevailing, XEC, a recombinant variant of KS.1.1/KP.3.3, shows a strong potential to become the next dominant strain, demonstrating a rapid expansion across multiple countries in Europe and North America. Compared to KP.3, XEC carries two additional Spike mutations, F59S and T22N (Figure A). Interestingly, both S31del of KP.3.1.1 and T22N of XEC introduce potential glycosylation on the N-terminal domain (NTD). Consequently, there is an imperative need to characterize the antigenicity and infectivity of KP.3.1.1 and XEC, especially focusing on the impact of NTD glycosylation.

**Figure.**
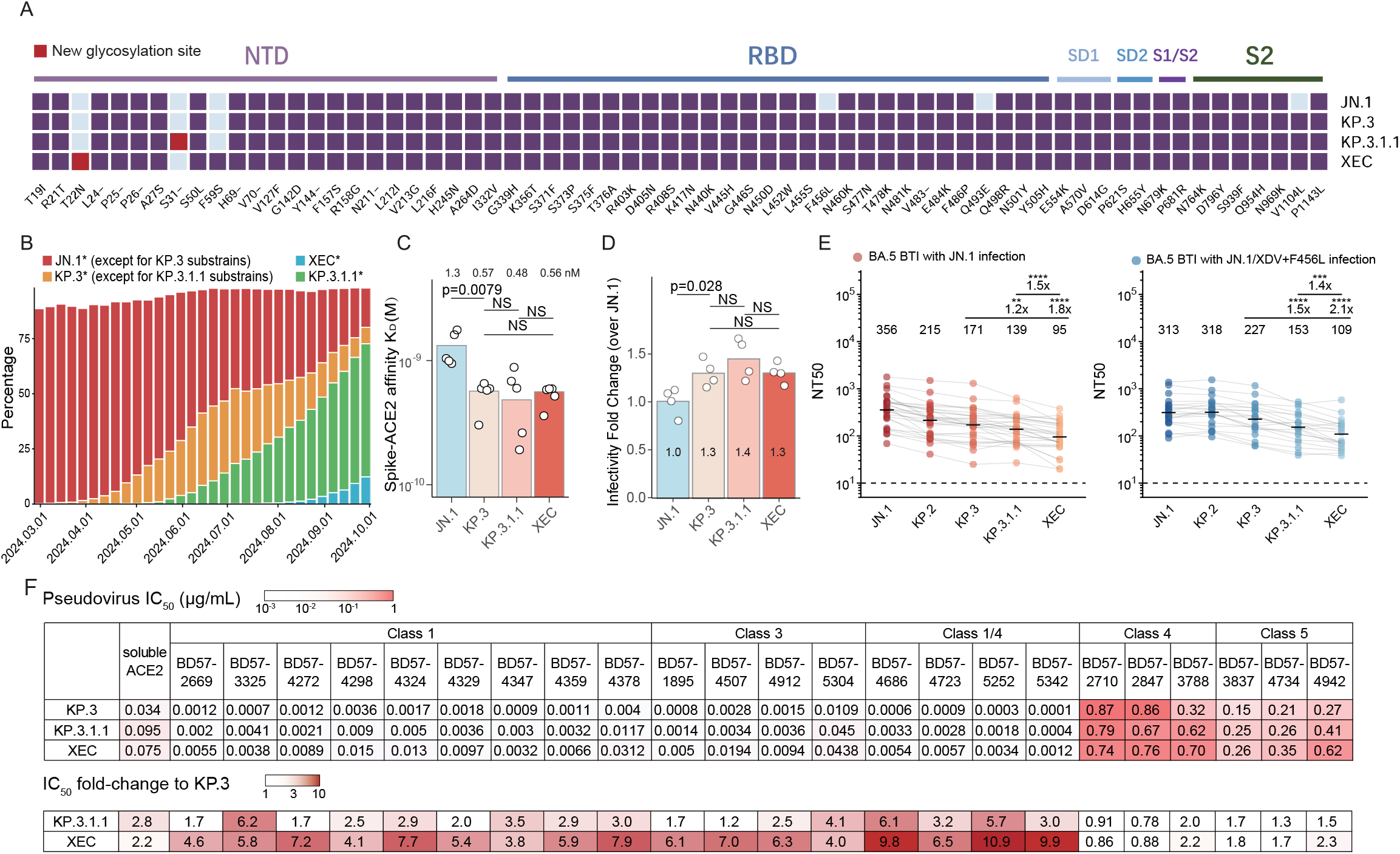
Infectivity and antibody evasion charaterization of KP.3.1.1 and XEC. (A) Spike protein mutations in JN.1, KP.3, KP.3.1.1, and XEC. Purple squares indicate mutations present in each variant, red squares denote new glycosylation mutations introduced by KP.3.1.1 and XEC, while sky-blue squares indicate the absence of mutations. (B) The percentage of circulating variants sequences globally from March 2024 to September 2024, including JN.1 (original JN.1 and its subvariants, excluding KP.3), KP.3 (original KP.3 and its subvariants, excluding KP.3.1.1), KP.3.1.1, and XEC. Data were collected from covSPECTRUM. (C) The binding affinity of JN.1, KP.3, KP.3.1.1, and XEC spike proteins to hACE2, determined by SPR. Each circle indicates a replicate. KD values (nM) are displayed above the bars. Two-tailed Wilcoxon signed-rank tests were performed to calculate p-values. (D) Relative infectivity of KP.3, KP.3.1.1, XEC compared with JN.1. The pseudovirus infectivity was tested using VSV pseudoviruses and Vero cells. Mean values are indicated on each bar. Two-tailed Wilcoxon signed-rank tests were used to calculate the p-values. (E) The 50% neutralization titers (NT50) of convalescent plasma from individuals reinfected with JN.1 after BA.5/BF.7 BTI (n = 29) and those reinfected with JN.1/XDV + F456L after BA.5/BF.7 BTI (n = 21). Geometric mean titers (GMT) are labeled above each group, with fold changes and statistical significance indicated above the GMT labels. Paired samples were analyzed using a two-tailed Wilcoxon signed-rank test. **p<0.01, ***p<0.001, ****p<0.0001. (F) The 50% inhibitory concentration (IC50, μg/mL) of a panel of monoclonal neutralizing antibodies targeting RBD epitopes against KP.3, KP.3.1.1, and XEC mutants. The table below presents the IC50 fold-change of KP.3.1.1 and XEC relative to KP.3. The antibody epitope groups are annotated above.

We first employed surface plasmon resonance (SPR) to assess the binding affinity between the Spike and human ACE2 (hACE2) of JN.1, KP.3, KP.3.1.1, and XEC. Consistent with previous findings, KP.3, KP.3.1.1, and XEC all exhibited an increase in ACE2-Spike binding affinity compared to JN.1^1,2,9^. However, we did not observe a significant change in the receptor binding of KP.3.1.1 and XEC relative to KP.3, despite the NTD glycosylation mutations (Figure C). Then, we evaluated the infectivity of KP.3.1.1 and XEC using vesicular stomatitis virus (VSV)-based pseudoviruses in Vero Cells. We found that although the pseudovirus infectivity of KP.3, KP.3.1.1, and XEC increased compared to JN.1, no significant differences were observed among KP.3, KP.3.1.1, and XEC (Figure D), consistent with recent report^10^.

Next, we conducted pseudovirus-based neutralization assays to assess the plasma neutralization escape of KP.3.1.1 and XEC. Specifically, we utilized the plasma from two cohorts: one cohort included vaccinated participants who experienced breakthrough infections (BTIs) with BA.5/BF.7 followed by reinfection with JN.1 (n=29), while the other cohort comprised individuals reinfected with JN.1/XDV + F456L (n=21) (Table S1). The specific variant causing reinfections was inferred from their local prevalence (>90%) at the sampling timepoint (Figure S2). Of note, XDV shares the same Spike sequence as JN.1. Importantly, we found that KP.3.1.1 and XEC exhibited significantly enhanced immune evasion capabilities than KP.3 (Figure E). The NT50 values against KP.3.1.1 decreased by 1.2-fold and 1.5-fold compared to KP.3, while that against XEC decreased by 1.8-fold and 2.1-fold. Notably, compared to the JN.1 reinfection cohort, the JN.1/XDV + F456L group exhibited a significant plasma neutralization improvement only against KP.2, whereas no differences were observed against KP.3, KP.3.1.1, and XEC (Figure S3).

Since KP.3.1.1 and XEC do not exhibit additional RBD mutations than KP.3, and the fact that most neutralizing antibodies should target the RBD instead of NTD, we did not expect the substantial plasma neutralization evasion of KP.3.1.1 and XEC. We hypothesized that the NTD glycosylation could affect the neutralizing activity of RBD-targeting NAbs through allosteric effects. Indeed, compared to KP.3, KP.3.1.1 and XEC exhibited enhanced escape capabilities against NAbs targeting various epitopes on the RBD, especially those compete with ACE2^1^ (Figure F). The inhibition efficiency of soluble ACE2 also showed slight reduction. This indicates that the glycosylation mutations on the NTD of KP.3.1.1 and XEC could reduce the potency of RBD-targeting NAbs via allostery, given the mutations are located away from the corresponding epitopes.

In summary, we found that KP.3.1.1, and especially XEC, exhibited enhanced humoral immune evasion and RBD-targeting antibody escape capabilities, supporting XEC’s foreseeable dominance. However, the detailed mechanisms by which T22N and S31del enhance antibody escape from RBD-targeting antibodies remain unclear and require further exploration. This may involve kinetic changes in the conformational dynamics of the RBD induced by additional glycosylation in the NTD, or it could be due to the impact of glycosylation sites on membrane fusion efficiency. Further structural studies are necessary to elucidate the mechanism.

## Acknowledgments

We thank the scientific community for their ongoing surveillance of SARS-CoV-2 variants and all volunteers for their blood sample contributions. This project is financially supported by the Ministry of Science and Technology of China (2023YFC3041500; 2023YFC3043200), Changping Laboratory (2021A0201; 2021D0102), and the National Natural Science Foundation of China (32222030, 2023011477).

## Author Contributions

Y.C. designed and supervised the study. J.L. and Y.C. wrote the manuscript with inputs from all authors. J.L., F.J., W.S., and S.Y. performed sequence analysis and illustration. Y.Y. and Youchun W. constructed pseudoviruses. P.W., L.Y., and F.S. processed the plasma samples and performed the pseudovirus neutralization assays. F.J. and Y.C. analyzed the neutralization data.

## Supplementary Figures

**Figure S1.**
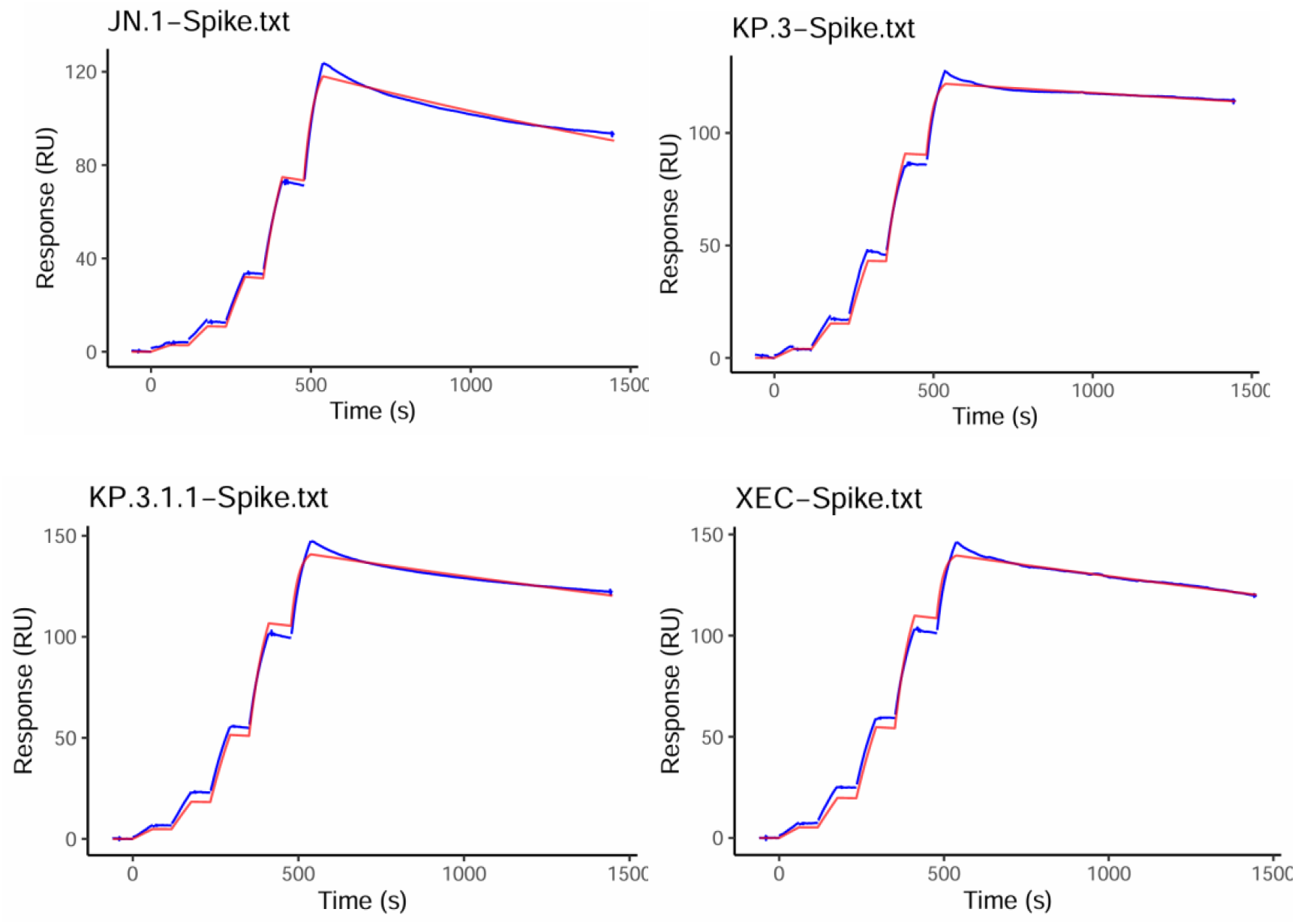
SPR sensorgrams of Spike-hACE2 binding. SPR sensorgrams to determine the hACE2-binding affinity of JN.1, KP.3, KP.3.1.1, and XEC Spike. Representative sensorgrams of replicates are shown. Kinetic parameters fitted by 1:1 binding model are annotated.

**Figure S2.**
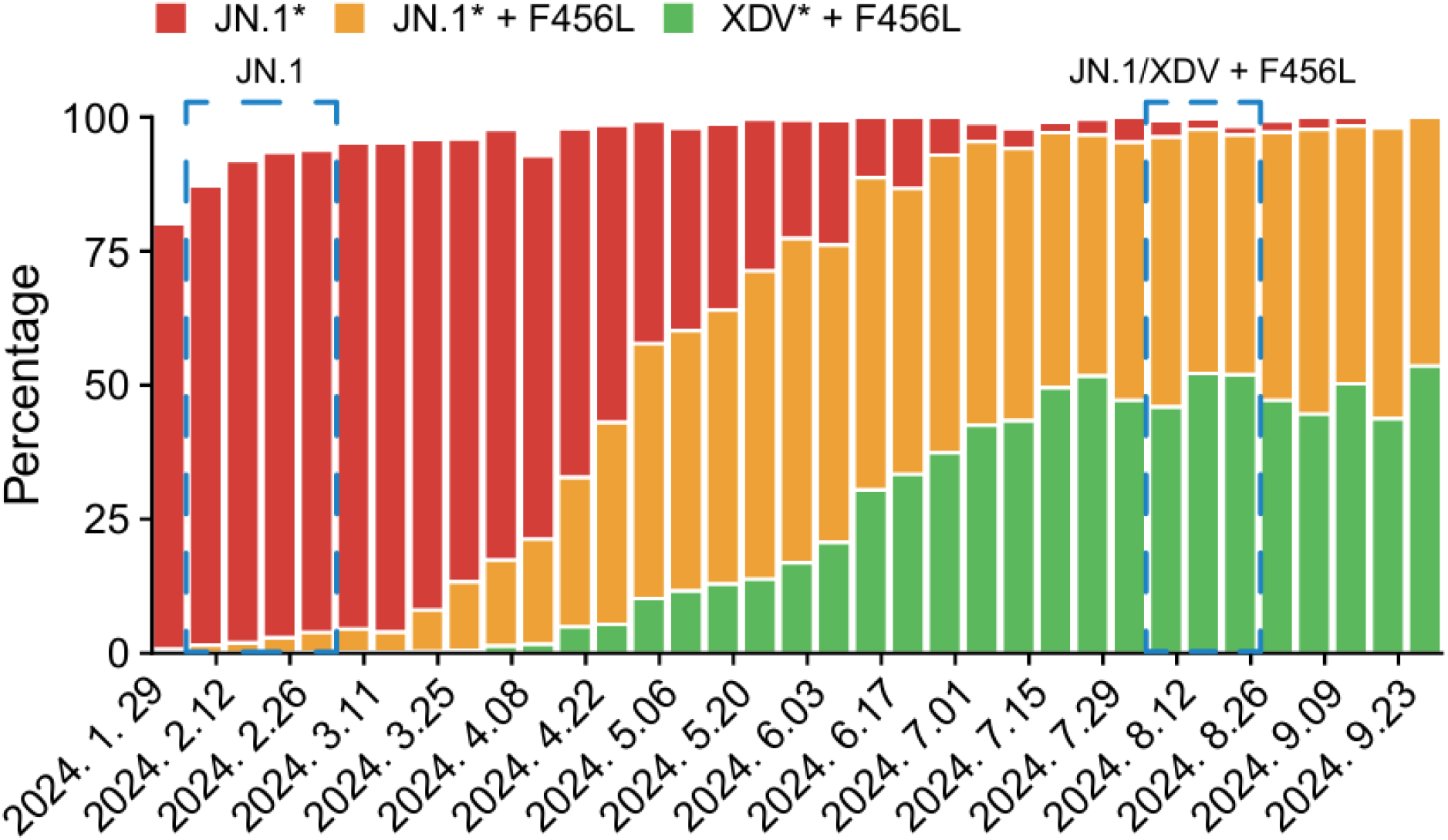
SARS-CoV-2 variant proportion in China during plasma sampling. The percentage of circulating variant sequences from February to October 2024 in China, including JN.1, JN.1 + F456L, and XDV + F456L. Data were collected from covSPECTRUM. The blue dashed line delineates the sampling time frame.

**Figure S3.**
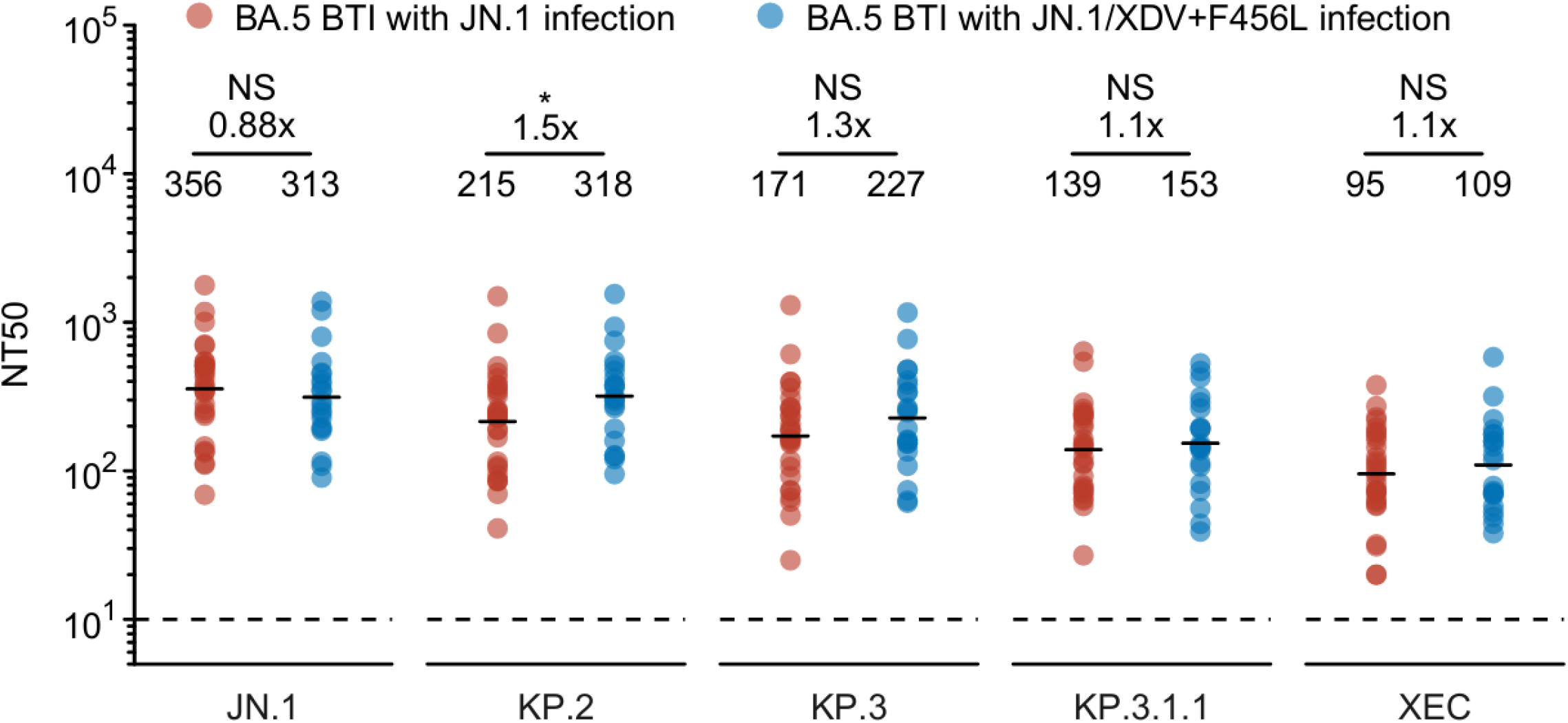
Neutralization comparison between JN.1 and JN.1/XDV + F456L. The 50% neutralization titers (NT50) of convalescent plasma from individuals reinfected with JN.1 after BA.5 BTI (n = 29) and those reinfected with JN.1/XDV + F456L after BA.5 BTI (n = 21) against SARS-CoV-2 variants JN.1, KP.2, KP.3, KP.3.1.1, and XEC. Geometric mean titers (GMT) are labeled above each group, with fold changes and statistical significance indicated above the GMT labels. Paired samples were analyzed using a two-tailed Wilcoxon signed-rank test. *p<0.05.

## Methods

### Patient recruitment and plasma isolation

Blood samples were collected from individuals who experienced reinfection with the SARS-CoV-2 Omicron BTI variant. This collection adhered to a research protocol approved by Beijing Ditan Hospital, affiliated with Capital Medical University (Ethics Committee Archiving No. LL-2021-024-02), the Tianjin Municipal Health Commission, and the Ethics Committee of Tianjin First Central Hospital (Ethics Committee Archiving No. 2022N045KY). Before sample collection, all participants provided written informed consent for the collection, storage, and exclusive research use of their blood samples, including the publication of related findings.

The patients’ initial infections occurred in December 2022 in Beijing and Tianjin, involving BA.5/BF.7 variants. As previously described, from December 1, 2022, to February 1, 2023, over 98% of sequenced samples were identified as BA.5* (excluding BQ*), predominantly comprising the subtypes BA.5.2.48 and BF.7.14, which represented the BA.5/BF.7 variants during this period. Patients in the JN.1 reinfection cohort experienced their first infection in December 2022 after receiving two or three doses of the COVID-19 vaccine, with their second infection occurring between February and March 2024. Similarly, patients in the JN.1/XDV + F456L reinfection cohort experienced their second infection from July to August 2024, following an initial breakthrough infection. From July 1, 2024, to September 1, 2024, over 97% of sequenced samples were confirmed as JN.1 + F456 or XDV + F456L (Figure S2). Confirmation of these infections was conducted using polymerase chain reaction (PCR) or antigen assays.

Following collection, whole blood samples were diluted 1:1 with PBS containing 2% FBS and subjected to gradient centrifugation using Ficoll (Cytiva, 17-1440-03). The resulting plasma samples were collected, aliquoted, and stored at -20°C or lower, undergoing heat inactivation before further experimental use.

### Pseudovirus neutralization assay

Utilizing the vesicular stomatitis virus (VSV) pseudovirus packaging system, we generated a pseudovirus of the SARS-CoV-2 variant spike. The G*ΔG-VSV virus (VSV G pseudotyped virus, Kerafast) was introduced into the cell culture supernatant. The pcDNA3.1 vector containing a mammalian codon-optimized spike gene was then transfected into 293T cells (American Type Culture Collection [ATCC], CRL-3216). Following cell culture, the pseudovirus in the supernatant was harvested, filtered, aliquoted, and stored at −80°C for future experiments. Plasma samples or antibodies were serially diluted in culture media, mixed with the pseudovirus, and incubated for 1 hour at 37°C in a 5% CO2 environment. Subsequently, digested Huh-7 cells (Japanese Collection of Research Bioresources [JCRB], 0403) were added to the antibody-virus mixture. After a 24-hour incubation period, the supernatant was discarded. D-luciferin reagent (PerkinElmer, 6066769) was then added and incubated in the dark for 2 minutes. Cells were lysed and transferred to detection plates. A microplate spectrophotometer (PerkinElmer, HH3400) was used to measure luminescence, and a four-parameter logistic regression model was applied to calculate the IC50 values.

### Surface Plasmon Resonance

SPR analyses were conducted on the Spike protein of the JN.1, KP.3, KP.3.1.1, and XEC, using a Biacore 8K system (Cytiva). Human ACE2, which was fused with an Fc tag, was immobilized onto Protein A sensor chips (Cytiva). Following serial dilutions of the purified Spike protein samples (6.25, 12.5, 25, 50, and 100 nM), the variants were injected onto the sensor chips. Data acquisition was performed using the Biacore 8K Evaluation Software 3.0 (Cytiva) at room temperature, and the resulting raw data were analyzed using a 1:1 binding model with the same software. Each variant was tested in two or three independent replicates to ensure reliability.

